# Cancer cells with alternative lengthening of telomeres do not display a general hypersensitivity to ATR inhibition

**DOI:** 10.1101/053280

**Authors:** Katharina I. Deeg, Inn Chung, Caroline Bauer, Karsten Rippe

**Author notes:** Correspondence should be addressed to K.R.

## Abstract

Telomere maintenance is a hallmark of cancer as it provides cancer cells with cellular immortality. A significant fraction of tumors uses the alternative lengthening of telomeres (ALT) pathway to elongate their telomeres and to gain an unlimited proliferation potential. Since the ALT pathway is unique to cancer cells, it represents a potentially valuable, currently unexploited target for anticancer therapies. Recently, it was proposed that ALT renders cells hypersensitive to ataxia telangiectasia-and RAD3-related (ATR) protein inhibitors (Flynn et al., Science 347, 273). Here, we measured the response of various ALT or telomerase positive cell lines to the ATR inhibitor VE-821. In addition, we compared the effect of the inhibitor on cell viability in an isogenic cell line, in which ALT was active or suppressed. In these experiments a general ATR inhibitor sensitivity of cells with ALT could not be confirmed. We rather propose that the observed variations in sensitivity reflect differences between cell lines that are unrelated to ALT.

## Introduction

Cancer cells need to maintain their telomeres to avoid cellular senescence and apoptosis induced by the replicative shortening of chromosome ends. Frequently, expression of TERT, the protein subunit of telomerase, is reactivated to extend the telomeres. In addition a significant fraction of tumors elongates the telomeres by an alternative lengthening of telomeres (ALT) pathway that operates via DNA repair and recombination processes as reviewed previously (Conomos et al., 2013; Pickett and Reddel,2015). ALT is not known to occur in healthy cells and thus represents a unique feature of cancer cells that could be targeted with specific drugs.

A recent study investigated telomerase positive and ALT positive osteosarcoma cell lines and glioma stem cells and reported that cells that employ the ALT pathway are hypersensitive to the inhibition of the protein kinase ataxia telangiectasia-and RAD3-related protein (ATR), one of the two main DNA damage checkpoint-activating kinases in human cells (Flynn et al., 2015).The authors concluded that treatment with the ATR inhibitor VE-821 selectively kills ALT cells within 6 days. They proposed that the immediate cell death induced by ATR inhibition in ALT cells is caused by an accumulation of DNA damage,aberrant anaphase chromosome segregation and increased micronuclei formation. Yet, how ATR inhibition elicits these effects specifically in ALT cells to affect the short-term cell viability remains elusive. Previously, several studies have demonstrated that inhibiting telomerase or ALT will induce senescence or cell death, e.g.(Asai et al., 2003;) (Jiang et al., 2005; (Potts and Yu, 2007; Shay, 2016). However, in the latter work many population doublings over weeks and months were required to elicit this type of response, in line with the view that no significant effects on cell viability due to telomere shortening are expected to occur within a few days.

Here, we recapitulated the cell viability and FACS experiments by Flynn et al.,. using various ALT or telomerase positive cell lines. Additionally, we investigated whether suppression of ALT activity affects cell viability upon treatment with ATR inhibitor. In our study no general hypersensitivity of ALT positive cells towards ATR inhibitors was observed.

## Materials and Methods

### Cell culture

Validated U2OS, HeLa, CAL72, HCT116 and SAOS2 cells were obtained from the German Collection of Microorganisms and Cell Cultures (DSMZ, Braunschweig, Germany). The MG63 cell line was purchased from CLS Cell Lines Service (Eppelheim, Germany). U2OS, HeLa and CAL72 cells were maintained in DMEM supplemented with 10% FCS, 2 mM L-glutamine, 1% antibiotics. For CAL72, 1X ITS liquid media supplement (Sigma-Aldrich) was added to this medium. The other three cell lines studied were cultured in 90% McCoy’s 5A supplemented with 10% FCS, 2 mM L-glutamine and 1% antibiotics (HCT116), in 85% McCoy’s 5A supplemented with 20% FCS (SAOS2) or in DMEM/Ham’s F12 medium supplemented with 5% FCS, 2 mM L-glutamine and 1% antibiotics (MG63).

### Cell viability assays

For cell viability assays, cells were seeded in triplicate in 96-well plates and incubated overnight. Different cell numbers were seeded to determine the cell number needed to obtain 70–90% confluency of the control sample after 6 days. Optimal starting cell numbers for U2OS, HeLa, HCT116 and MG63 were 500 cells, and for CAL72 and SAOS2 1500 cells. The following day cells were either treated with DMSO (control) or with increasing concentrations (0.5, 1, 2, 4, 8, 16 μ) of the ATR inhibitor VE-821 (Selleckchem) dissolved in DMSO. Cells were incubated for 6 days without medium change and cell viability was analyzed using CellTiter Glo (Promega) and a TECAN Infinite M200 plate reader according to the manufacturers’ instructions.

### FACS analysis of cell death

For analysis of cell death, cells were seeded in T25 flasks. Cell numbers were adjusted for each cell line to account for varying proliferation rates (1× 10^5^ CAL72 cells, 1.4 × 10^5^ SAOS cells and 0.8 × 10^5^ cells for HeLa, HCT116 and U2OS). The following day each cell line was either treated with 3 μ VE-821 (Selleckchem) or with the same volume of DMSO for the control samples. Cells were incubated for 6 days without medium change. Cells, including dead cells, were collected by trypsin and total cell numbers were determined using the LUNA cell counter (Biozym). Cells were resuspended in FACS binding buffer (10 mM HEPES, 2.5 mM CaCl_2_, 140 mM NaCl) at a final concentration of 2×10^6^ cells/ml, stained with FITC annexin V (BioLegend) and propidium iodide (Miltenyi Biotec) according to the manufacturers’ instructions, and analyzed by flow cytometry on a FACS Canto II (BD Biosciences). The fraction of apoptotic cells, characterized as annexin V positive, was quantified using the FACS Diva software. The percentage of induced cell death d_ind_ was calculated as

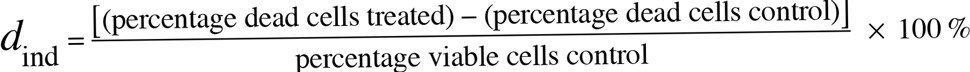

### Generation and analysis of a U2OS cell line with inducible ATRX expression

PEGFP-C2-ATRX-HA was kindly provided by David Picketts (University of Ottawa, Canada) (Berube et al., 2008) ATRX-HA was amplified and cloned into the pTRE3G-ZSGreen1 vector (Clontech, USA) to construct pTRE3G-ZSGreen1-ATRX-HA for doxycycline inducible expression of ATRX. Plasmids that contained the ATRX cDNA were propagated in the *dam/dcm*-negative bacteria strain JM110 (Agilent Technologies) to avoid transposon insertions. A stable U2OS^ATRX^ cell line was generated by transfecting U2OS cells with pCMV-Tet3G (Clontech, USA) and pTRE3G-ZSGreen1-ATRX-HA and subsequent selection with G418 (1 mg/ml). U2OS^ATRX^ cells were cultured under the same conditions as the parental U2OS cell line except that doxycycline-free FCS was used. For induction of ATRX expression 1 μg/ml doxycycline was added to the medium. ALT activity after ATRX expression was evaluated by the C-circle assay and the number of ALT-associated PML bodies (APBs) as described previously (Osterwald et al., 2015). U2OS^ATRX^ (-) and U2OS^atrx^ (+) cells were cultured in the absence or presence of 1 μg/ml doxycycline, respectively, for at least 7 days before treatment with ATR inhibitor. For the cell viability assay 1000 cells of U2OS^ATRX^ (-) and U2OS^ATRX^ (+) were initially seeded and treated with inhibitor as described above.

## Results and Discussion

To evaluate the sensitivity of cell lines to ATR inhibition, we compared the human telomerase positive (“TEL”) cell lines HeLa, HCT116 and MG63 with the ALT cell lines U2OS, CAL72 and SAOS2. Cell viability measurements after treatment with different ATR inhibitor concentrations were conducted using the CellTiter Glo assay (Promega) and by fluorescence activated cell-sorting (FACS) analysis with annexin V staining. The experimental methods and inhibitor treatment conditions of 6 days incubation without medium change were those used previously by Flynn et al., who also studied the MG63, U2OS, CAL72 and SAOS2 osteosarcoma cell lines (Flynn et al., 2015).

### Cell viability analysis with the CellTiter Glo assay

In order to measure cell viability in response to VE-821 we first examined the HCT116 TEL cell line (Fig. 1A) in reference to the original paper that characterized the VE-821 inhibitor (Flynn et al., 2015 Reaper et al., 2011). It is noted that the CellTiter Glo assay measures changes in the number of viable cells after inhibitor treatment relative to a control sample. Whether the resulting differences originate from a reduced growth rate or from an increase in cell death cannot be distinguished in this assay. The half-maximal inhibitory concentration (IC^50^) for the HCT116 cell line in our experiments was 1-2 μM and similar to that reported previously (see Fig. S4D in (Reaper et al., 2011)). In the course of this study we noted that the results were dependent on the amount of cells initially seeded (Fig. 1A). Seeding higher cell numbers shifted the dose response curve to elevated inhibitor concentrations. Consequently, the cells appeared less sensitive to the inhibitor. The same dependence was observed for the U2OS ALT cell line (Fig. 1B). This behavior could be the result of several factors: Cells may be less sensitive to ATR inhibitor treatment at higher starting cell numbers as they have more cell-to-cell contacts and are thus more resistant to stress caused by the treatment. Additionally, cells can appear less sensitive to the ATR inhibitor when starting with higher cell numbers because the proliferating control cells reach confluency before the treatment ends after 6 days. Thus, the control cells would have already stopped dividing further before the end point of the experiment is reached. In contrast, cells growing slower upon treatment may not reach confluency within 6 days and continue to proliferate during the complete observation period, albeit at a lower rate. Since values are normalized to the control, this would make the ratio of viable cells in treated versus control samples dependent on the density of seeded cells, the proliferation rate and the observation time.

**Figure 1.**
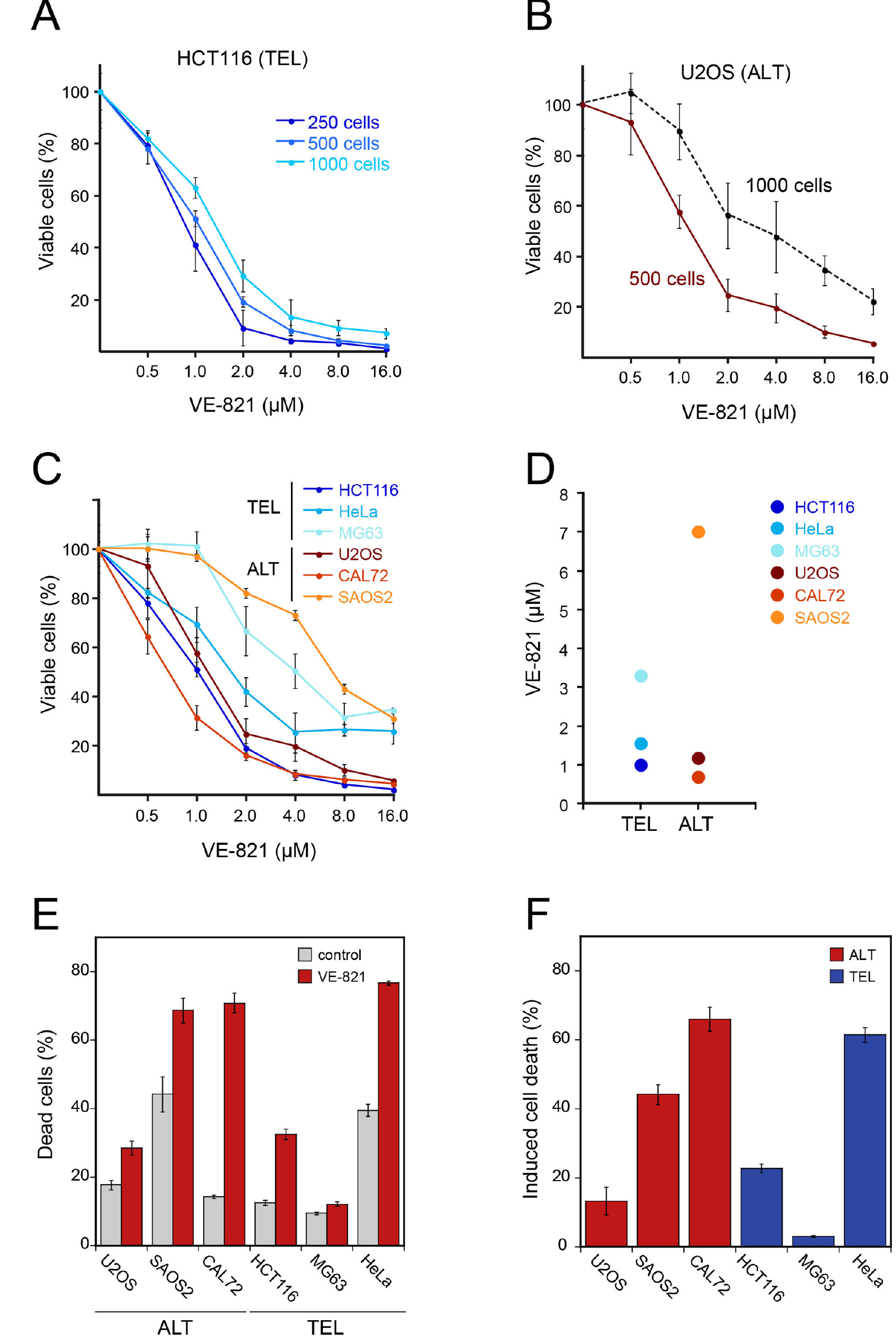
**Cell viability of ALT and telomerase positive (TEL) cell lines upon treatment with the ATR inhibitor VE-821** (**A**) Different cell numbers of HCT116 (TEL) cells were seeded in a 96-well plate and treated with increasing concentrations of the ATR inhibitor VE-821 for 6 days to determine the influence of the starting cell number on the cell viability assay. Cell viability was analyzed using CellTiter Glo and is depicted as percentage of control (*n* = 3). (**B**) Same as panel *B* but for the U2OS ALT cell line. (**C**) Analysis of cell viability for different cell lines with starting cell numbers that led to 70-90% confluency after 6 days in the absence of inhibitor (*n* = 3). Cell numbers used for seeding were 500 cells/well for U2OS, HeLa, HCT116 and MG63 and 1500 cells/well for the CAL72 and SAOS2 cell lines. Error bars represent SD of triplicate experiments. (**D**) IC_50_ concentrations determined from cell viability curves shown in panel *C* by fitting a sigmoidal curve. (**E**) FACS analysis of cell death in VE-821 treated ALT and TEL cell lines. Fraction of dead cells in ALT and TEL cell lines treated with DMSO (control) or 3 μM VE-821 for 6 days. Dead cells were determined by FACS analysis of annexin V stained cells. Error bars represent SEM (*n*≥2). (**F**)Percentage induced cell death determined from FACS analysis as in panel E. Induced cell death was calculated as described in the Materials and Methods section.

### ATR inhibitor sensitivity of different cell lines

For comparing ALT and TEL cell lines we therefore selected a starting cell number for each cell line that led to 70–90% confluency after 6 days in the absence of the inhibitor. The optimized starting cell numbers used for U2OS, HeLa, HCT116 and MG63 were 500 cells, and for the slower growing CAL72 and SAOS2 cell lines 1500 cells per well of a 96-well plate. Using these seeding cell numbers we found no hypersensitivity of ALT cell lines in response to the VE–821 ATR inhibitor (Fig. 1C). Instead, the sensitivity varied between cell lines irrespective of the telomere maintenance mechanism. While the HCT116 TEL and the CAL72 ALT cell lines were sensitive to ATR inhibition at VE–821 concentrations of 1–2 μM, the MG63 TEL and the SAOS2 ALT cell lines showed a stronger reduction of viable cells only at higher inhibitor concentration. The IC_50_ value measured ranged from 0.9 to 3.3 μM for the TEL cell lines and from 0.7 to 7 μM for the ALT cell lines (Fig. 1D). Thus, the IC_50_ values of the two groups were not systematically different, but rather showed a high variation within each group.

Next we quantified dead cells by FACS analysis of annexin V stained telomerase and ALT positive cells treated with 3 μM VE-821 for 6 days (Fig. 1E, F). Measurements of the SAOS2 ALT cell line and the HeLa TEL cell line revealed a fraction of 40% dead cells already in the control samples in the absence of inhibitor (Fig. 1E). This indicates that these cells are more sensitive to 6 days of culturing without medium change. In comparison to the control, the CAL72 ALT cell line displayed the highest sensitivity to ATR inhibitor treatment in this assay, while the MG63 TEL cells were mostly insensitive as reflected by the calculated percentages of induced cell death in relation to the control (Fig. 1F). However, treatment with the ATR inhibitor induced a higher percentage of cell death in the HeLa and HCT116 TEL cell lines compared to the U2OS ALT cells. Thus, we did not observe a selective killing of ALT cells by ATR inhibition in these experiments.

### ATR inhibitor sensitivity in dependence of ALT activity

To evaluate the effect of ALT activity on ATR sensitivity in an isogenic cell line and independent of the above mentioned confounding factors we exploited the recent finding that the a-thalassemia mental retardation X-linked (ATRX) chromatin remodeling protein acts as a suppressor of ALT (Clynes et al., 2015; Napier et al., 2015). Accordingly, a cell line for the inducible expression of ectopic ATRX in the U2OS cell line that intrinsically harbors large deletions in the *ATRX* gene was established. In the resulting U2OS^ATRX^ cell line HA-tagged ATRX protein is produced upon induction as confirmed by western blot using an ATRX-and an HA-specific antibody (Fig. 2A). Upon induction of ATRX protein expression, ALT activity became progressively reduced as apparent from monitoring two characteristic ALT markers: single-stranded circular C-rich extrachromosomal telomere repeats (C-circles) as well as PML-telomere colocalizations, termed ALT-associated PML bodies (APBs) (Yeager et al., 1999; Henson et al., 2009). After 7 days of ATRX expression, the number of APBs was at the background level observed for TEL cell lines(Fig. 2B)and C-circles were almost undetectable (Fig. 2C) indicating a complete inhibition of ALT activity. Next, we compared the ATR inhibitor sensitivity of U2OS^ATRX^ (+) cells that had ALT silenced due to ATRX induction to the same U2OS^ATRX^ (−) uninduced cell line with an active ALT pathway. The dose-response curves for the two cell samples were identical as determined with the CellTiter Glo assay (Fig. 2D). Thus, silencing ALT activity via ectopic expression of ATRX did not affect ATR inhibitor sensitivity of the cells.

**Figure 2.**
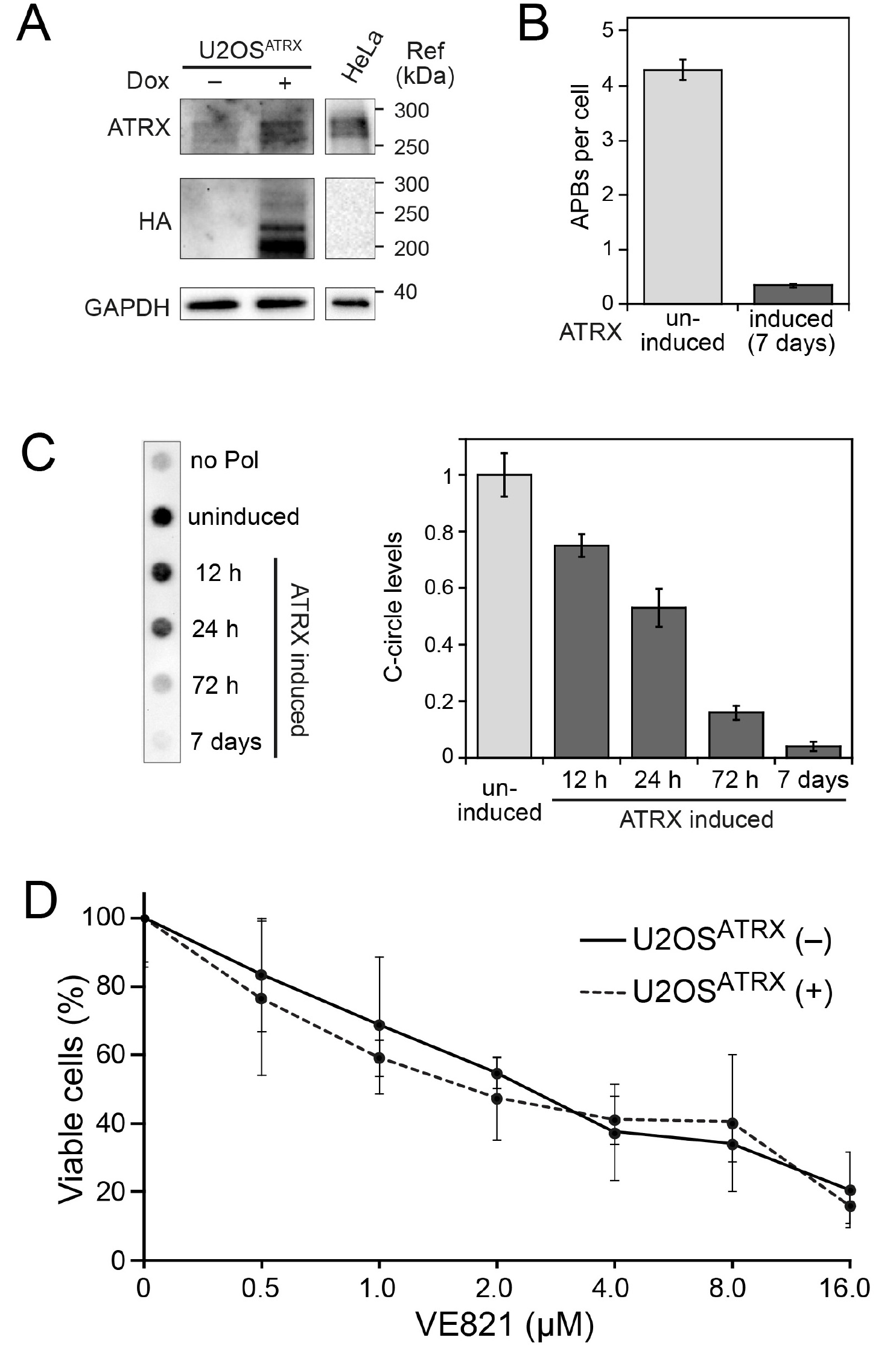
**ALT features and sensitivity to ATR inhibitor treatment upon ectopic expression of ATRX in ATRX-deficient U2OS cells.** (**A**) Western blot showing the expression of HA-tagged ATRX in the generated U2OS^ATRX^ cell clone upon doxycycline induction for 48 h and a HeLa reference. In addition to the full-length ATRX band at about 260 kDa, shorter ATRX variants between 200 and 220 kDa were also detected in the HA blot that might correspond to degraded or alternatively spliced products of ATRX, as described previously (Berube et al., 2000); (Mitson et al., 2011.) (**B**)Average number of APBs per cell in uninduced and for 7 days induced U2OS^ATRX^ cells (*n* = 350) analyzed by 3D confocal image analysis of PML IF and telomere FISH stainings as described previously (Osterwald et al., 2015.) (**C**)C-circle assay as a marker of ALT activity in uninduced and induced U2OS^ATRX^ cells. Samples without polymerase (no pol) and uninduced U2OS^ATRX^ were included as controls. The bar blot shows a quantification of C-circle levels in uninduced and induced U2OS^ATRX^ cells from three experiments. (**D**)ATR inhibitor sensitivity in dependence of ALT activity in the U2OS^ATRX^ cell line. ATRX induced U2OS^ATRX^ (+) and uninduced U2OS^ATRX^ (—) cells were analyzed using the CellTiter Glo assay in the presence of increasing concentrations of the ATR inhibitor VE-821 for 6 days. No changes in ATR inhibitor sensitivity were observed when ALT was silenced by ATRX expression.

## Concluding remarks

ATR and the protein kinase ataxia telangiectasia mutated (ATM) are the two main DNA damage checkpoint-activating kinases in human cell. Consistent with the view that replication stress and misguided DNA repair synthesis are crucial features of ALT it was found that inhibition of ATR or ATM decreases ALT activity (Nabetani et al., 2004; O’sullivan et al., 2014; Flynn et al., 2015;(Osterwald et al., 2015.). However, except for the Flynn et al. study, no immediate ALT specific effects after ATR and/or ATM inhibition on cell viability and proliferation on the time scale of several days have been reported.

In our comparison of different cell lines we identified a number of factors that affected the apparent sensitivity towards the VE–821 ATR inhibitor but were unrelated to ALT (Fig. 1). We found that the initial cell number seeded as well as intrinsic differences in proliferation rates between cell lines have to be considered when interpreting measurements of cell viability assays. Furthermore, we note that a reduction in cell viability detected in the CellTiter Glo assay does not allow discrimination between a reduced growth rate versus an increase of cell death. As an additional parameter the experimental procedure of treating cells with inhibitor for 6 days without changing medium might introduce a bias against cells that are sensitive to factors that accumulate in the medium or become depleted upon prolonged cultivation. Finally, differences in the genetic background may lead to ATR inhibitor sensitivity independent of ALT. For example, the telomerase positive HCT116 colon cancer cell line used here harbors a mutation in *MRE11*, which impairs binding to NBS1 and Rad50 and suppresses ATM activation in response to replication stress (Giannini et al., 2002; Wen et al., 2008). This may account for its relatively high sensitivity towards ATR inhibition in terms of cell viability independent of its telomere maintenance mechanism.

In addition to the effects of the above mentioned factors it would still be conceivable that the presence of ALT contributes to an increased sensitivity to ATR inhibition. To test whether this is the case we compared a U2OS cell line in which ALT was active with the same cell line that had ALT silenced by inducing ectopic ATRX expression. In these experiments, the ATR inhibitor sensitivity was not changed when the ALT pathway was rendered inactive (Fig. 2). Thus, we conclude that cells that employ ALT to maintain their telomeres are not generally more sensitive to ATR inhibition than telomerase positive cells on the time scale of days. Rather we suggest that, as described above, the cell line specific genetic background and additional factors exist that are responsible for the observed effect of ATR inhibition.

Our results indicate that ATR inhibition alone will not be sufficient to target tumors in which ALT is active. Nevertheless, we share the view that the misguided DNA repair and recombination mechanism active in ALT provides unique novel options for anti-cancer therapies. In this context, the recurrent inactivation of the ATRX tumor suppressor protein in ALT cancer samples could be exploited (Watson et al., 2015).As inactive ATRX is associated with ALT specific tumor features, it could for example be targeted by synthetic lethality approaches. In support of this conclusion, it has been recently shown that ATRX deficiency impairs non-homologous end joining and increases sensitivity to DNA-damaging agents in a glioma mouse model (Koschmann et al., 2016). A systematic further investigation of this relation appears to be promising for exploiting ALT associated cellular deregulation in personalized cancer therapies.

## Acknowledgements

This work was supported within project CancerTelSys (01ZX1302) in the e:Med program of the German Federal Ministry of Education and Research (BMBF).

